# Tackling the research capacity challenge in Africa: An overview of African-led approaches to strengthen research capacity

**DOI:** 10.1101/518498

**Authors:** Lem Ngongalah, Ngwa Niba Rawlings, Emerson Wepngong, James Musisi, Claude Ngwayu, Sharon Mumah

## Abstract

**Background:** Improved capacity for research is a valuable and sustainable means of advancing health and development in Africa. Local leadership in research capacity strengthening is important for developing contextually appropriate programs that increase locally-driven research, and improve Africa’s ability to adapt and use scientific knowledge. This study provides an overview of African organisations that aim to strengthen research capacity in Africa, and the major initiatives or approaches being used for this purpose.

**Methods:** A desk review of grey and published literature on research capacity strengthening in Africa was conducted, in addition to panel discussions on the determinants of research capacity in Africa. Data was analysed through thematic analysis and a framework developed by the Collaboration for Research Excellence in Africa (CORE Africa).

**Results:** 11 organisations were identified, spread across South, Central, East and West Africa. The main approaches to improving research capacity were: providing opportunities for academic research and research training. Initiatives to provide research equipment, funding and facilitate research use for policy-making were limited; while strategies to increase research awareness, promote collaboration, and provide guidance and incentives for research were lacking. Most organisations had programs for researchers and academics, with none targeting funders or the general public.

**Conclusion:** Local leadership is essential for improving research capacity in Africa. In addition to providing adequate support to academics and researchers, initiatives that help revitalize the education system in Africa, promote collaboration and engage funders and the general public will be helpful for strengthening research capacity in Africa.

## 1. Introduction

Building capacity to produce and publish research is an important prerequisite for addressing existing and emerging health and development challenges in Africa. Research capacity is the ability to identify problems, set priorities, conduct sound scientific research, build sustainable institutions, and develop lasting solutions to key problems [1]. This definition embodies capacity at individual, institutional and national levels. The disease burden in Africa is significantly higher than that in other regions of the world [2]. Meanwhile, the technical and human capacity available to tackle these problems does not parallel the existing challenges [3, 4]. Poor research capacity significantly hinders the ability to build a local evidence base with which to inform policy and program development. While some African countries have seen progress in recent years, Africa continues to be under-represented in global research activities and output, at all levels of research [5].

The need to strengthen research capacity in Africa has long been recognised. Research capacity strengthening (RCS) refers to any efforts that increase the ability of individuals and institutions to undertake high-quality research, both individually and collectively, in an efficient and sustainable manner [6, 7]. This process has been identified as one of the most powerful, cost-effective, and sustainable means of advancing health and development in Africa [3]. Over the years, research knowledge and expertise from high-income countries (HICs) has helped identify preventive and therapeutic interventions for major causes of mortality in Africa, such as malaria and HIV/AIDS; and the development of health infrastructures and practices in Africa [8–10]. RCS efforts which have had a substantial impact in Africa have mostly been led by HIC institutions, and heavily dependent on continued foreign support [11]. Local leadership in RCS is important for developing contextually appropriate programs that help increase locally-driven research, and improve Africa’s ability to generate, adapt and use scientific knowledge. African researchers are best placed to identify challenges of their own nations and provide relevant evidence to policy-makers to inform decision-making.

Current literature on RCS in Africa is large and diverse, with varying ideologies and concepts. RCS is a dynamic and broad process requiring a combination of short and long-term strategies, which cannot be defined by any single model, framework or set of approaches. RCS initiatives and interventions in Africa are typically conceptualized around three levels of research: individual, organisational and institutional [12]. A number of recommendations have been proposed, such as: improving the research environment, supporting researchers and research institutions [7]; increasing collaboration for research in Africa [13]; increasing funding for postgraduate research [14]; improving communication about RCS work and investing in monitoring and evaluating capacity building [6]. However, the priorities and contributions of African-led organisations towards strengthening research capacity in Africa remain unclear.

The purpose of this report is to provide an overview of African organisations that aim to strengthen research capacity in Africa, and the major initiatives or approaches being used for this aim. This data is further analysed to identify gaps and opportunities that future RCS initiatives can build on. This study also aims to inform the RCS mission of the Collaboration for Research Excellence in Africa (CORE Africa). As part of its broader commitment to develop a solid and sustainable research workforce in Africa, CORE Africa has recognised the importance of strengthening research capacity. This report would help identify areas where CORE Africa can best contribute, and opportunities for collaboration and/or complementarity. The findings presented herein would also inform plans from other stakeholders working towards the realisation of sustainable, locally-led research in Africa.

## 2. Methods

This report is a distillation of different pieces of information and data, derived from various online sources including organisational websites, published literature, conference reports, and analytical or empirical documents. The study adopted a desk review approach to identify grey and published data on RCS in Africa. This method was used based on the fact that the type of information and data sought for this study would not generally be captured in a traditional review of published literature. The objectives of the study were to: a) identify African-led and Africa-based organisations that aim to strengthen research capacity in Africa; b) identify the RCS-related aims and objectives of these organisations; c) identify the methods used to achieve their aims/objectives; d) identify their target beneficiaries and e) identify collaboration activities with other African organisations.

### 2.1. Identification of RCS Organisations

A four-step search methodology was used. The initial search was done by reviewing a database of research institutions and organisations in Africa compiled by the institute of development studies (IDS). The IDS is an organisation that aims “to reduce inequalities, and accelerate sustainability by mobilising high quality research and knowledge that informs policy and practice” [15]. The list included universities, non-governmental organisations, networks and associations, bilateral agencies, philanthropic organisations, multilateral organisations and research councils.

The second step used to identify research organisations was by searching through the partners and collaborators of institutions identified in the first step. Additional organisations were identified through part of a survey previously administered by CORE Africa [16], where participants were asked to name any organisations they knew of, that provided research training in their respective countries. Our searches were then supplemented with a google search, using a variety of keywords including “research capacity”, “capacity building”, “research capacity strengthening”, “Africa”, “strengthening research capacity “improving research capacity” and “Sub-Saharan Africa”. Screening of all potentially-eligible organisations was done in duplicate and discrepancies were resolved through a third opinion.

### 2.2. Selection criteria

The eligibility criteria for this study included the following: organisations led or managed by Africans, organisations with main offices or headquarters in Africa, organisations focused on health-related research in Africa, organisations with aims or objectives that include building, improving or strengthening research capacity in Africa, and organisations that have programs, initiatives or activities related to these aims. Governments, universities, research groups and organisations which were specific to one area of research e.g. HIV, Malaria, etc were excluded.

### 2.3. Data analysis

The first phase of data analysis for this study used a thematic analysis approach, which is the process of identifying patterns or themes within qualitative data [17]. This was done by reading the collected information over and over to get familiar with the data, after which codes were generated for different aims and activities of each included organisation. The codes were then explored for similarities and differences, after which similar codes were clustered into themes. Resulting findings from these steps are described narratively.

In the second phase of analysis, a framework developed by CORE Africa was used to assess priority areas for the RCS organisations identified. This framework offers a simplistic approach to assess RCS activities in Africa, using relevant factors that influence research capacity. The framework was developed from information generated through the analysis of the literature, policy documents and empirical studies. The project team also held panel discussions with experienced researchers and other stakeholders on the determinants of research capacity, and how to integrate this knowledge into RCS initiatives in Africa. These findings were then translated into key factors and population groups that influence research capacity [18]. The basis of our analysis therefore, is that these factors should be addressed in the RCS process. Because every research capacity strengthening initiative occurs in a unique context, the factors outlined in this framework are intended to be informative, and to be taken on as a tool for further discussion by other RCS stakeholders.

## 3. Results

### 3.1. Description of RCS initiatives and organisations

The search resulted in a total of 338 initiatives/organisations, out of which 11 were eligible for inclusion in our study (table 1). These were spread over 11 countries in Sub-Saharan Africa - four in Southern Africa (South Africa, Malawi, Zambia and Mozambique); three each in East Africa (Uganda, Kenya, Tanzania), three in Central Africa (Cameroon, Democratic Republic of Congo, Rwanda) and one in West Africa (Nigeria) (figure 1). Web links for all included organisations are provided in Appendix A.

**Table 1:** RCS Organisations and initiatives included in the study.

| Organisation | Countries | RCS aims and objectives |
| --- | --- | --- |
| African Population and Health Research Centre (APHRC) | Kenya | To generate an African-led and African-owned body of evidence to inform decision making in Africa.<br>To strengthen the research capacity and skills of African scholars<br>To support evidence-based policy and decision making in Sub-Saharan Africa |
| African Institute for Development Policy (AFIDEP) | Kenya, Malawi | To strengthen the capacity of researchers, advocates, and policymakers in translating and using research and related forms of evidence |
| Research on Poverty Alleviation (REPOA) | Tanzania | To increase the number of researchers capable of undertaking policy-relevant quality research<br>To facilitate and inspire stakeholders to utilize research findings<br>To undertake, facilitate and encourage strategic research to influence policy |

| **Organisation** | **Countries** | **RCS aims and objectives** |
| --- | --- | --- |
| Information Training and Outreach Centre for Africa (ITOCA) | South Africa | To build capacity for scientist, researchers and information professionals on the use of electronic resources in Sub Saharan Africa (SSA) |
| The Africa Institute of South Africa (AISA) | South Africa | To provide training and produce research of the highest standard by working with the best researchers to contribute to development and knowledge creation for all Africa. |
| The Alliance for Accelerating Excellence in Science in Africa (AESA) | Kenya | To accelerate world-class research, foster innovation and promote scientific leadership in Africa |
| Southern Africa consortium for research excellence (SACORE) | South Africa | To establish a vibrant environment to support training and research in Southern Africa |
| Initiative to Strengthen Health Research Capacity in Africa (ISHReCA) | South Africa, Kenya | To promote the creation of self-sustaining pools of excellence capable of initiating and carrying out high quality health research in Africa. |
| The Consortium for Advanced Research Training in Africa (CARTA) | Uganda, Kenya, Nigeria, Tanzania, Malawi, Rwanda, South Africa | To strengthen research infrastructure and management capacity at African universities, and to support doctoral training through a model collaborative PhD program in population and public health that trains and retains the continents brightest minds. |
| Africa Centre of Excellence for Infectious Diseases of Humans and Animals in Eastern and Southern Africa (SACIDS-ACE) | Tanzania, Democratic Republic of Congo, Mozambique, South Africa, Zambia | To enhance Africa's research capacity by linking academics and research institutions |

| Organisation | Countries | RCS aims and objectives |
| --- | --- | --- |
| Clinical Research Education, Networking and Consultancy<br>(CRENC) | Cameroon | To promote excellence at each step of the research process, starting from evidence generation to its translation into clinical or public health policies for implementation |

**Figure 1:**
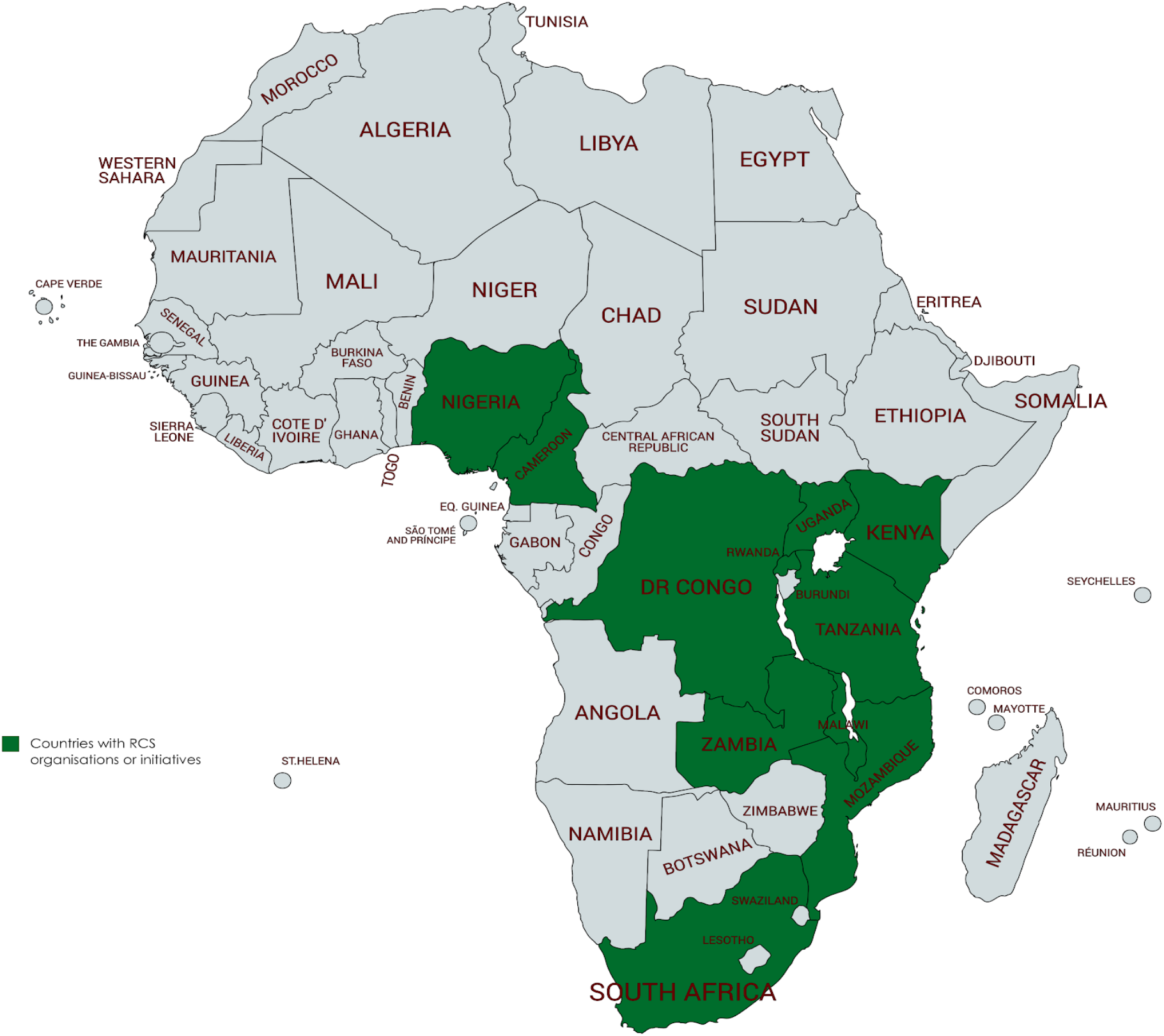
African countries where RCS organisations or initiatives were identified

### 3.2. Summary of RCS initiatives and activities

Six themes emerged from the analysis of RCS aims, initiatives and activities of the included organisations. These were: generating research evidence to influence research capacity, development of research skills, facilitating the uptake and use of research knowledge, providing funding and other resources for research, increasing research awareness and research collaboration.

#### 3.2.1. Generating research evidence to influence research capacity

Three of the identified organisations aim to generate an African-led and African-owned body of research knowledge to influence research capacity. These include APHRC, AESA and ISHRECA. APHRC conducts research on what works to make research and higher education systems sustainable in Africa. AESA functions as a think tank, ensuring an Africa-centred and Africa-relevant science and technology agenda on the continent. ISHRECA is an initiative that provides a platform for discussion of health research needs, and produces knowledge on how to improve health research capacity in Africa.

#### 3.2.2. Development of research skills

##### a) Providing research training opportunities

A majority (Nine out of eleven) of the identified organisations had programs that aim to develop or enhance research skills for African researchers. These were either academic research courses (e.g. Masters programs) or opportunities to develop practical research skills (table 2).

**Table 2:** Skill development programs offered by RCS organisations

| Type of skill development program offered | Organisation |
| --- | --- |
| MSc. programs, research training programs, internships, fellowships and short courses | APHRC, AFIDEP, REPOA, AISA, SACORE, CARTA, SACIDS-ACE, ITOCA, CRENC |
| Workshops, seminars | AFIDEP, ITOCA, CARTA, SACIDS-ACE |

##### b) Providing research guidance

Research guidance refers to any activities that enable less-experienced researchers to benefit from leadership, advice, supervision or counselling provided by more experienced researchers. The only form of guidance for research identified in this study was through mentorship. Providing research mentorship was one of the objectives of AESA – an initiative created by the African Academy of Sciences (AAS) and the New Partnership for Africa’s Development (NEPAD). However, attempts to access the AESA mentoring platform were futile, and this study was unable to obtain further information.

#### 3.2.3. Facilitating the use of research findings for policy-making

Two organisations were identified which aim to help bridge the gaps between research, policy and practice; in order to enable the creation of research-informed policies in Africa. AFIDEP does this by providing training on evidence synthesis, policy analysis, scenario building, forecasting and effective communication. REPOA provides training for research users on how to analyse and interpret research findings, and translate these into policy-related recommendations and interventions.

#### 3.2.4. Providing funding and other resources for research

##### a) Funding

This study identified four organisations with funding opportunities for research in Africa. ITOCA funds outreach programmes in information and communication technology (ICT) and related activities, and assists institutions in seeking funding to implement these programs. CARTA provides scholarships for PhD research. AESA awards grants for innovative solutions in Africa, and funds programs that address scientific quality, research training, mentorship, leadership, collaboration in science, research management, research environments and engagement with public and policy stakeholders. SACORE was an initiative consisting of three African universities and their affiliated research institutions, which provided fellowships and grants to researchers and academics from partner institutions [19].

##### b) Other resources and support for research

There were four organisations providing non-financial forms of support for research. These include: CRENC, SACIDS-ACE, AISA and ITOCA. CRENC provides educational support to postgraduate students on clinical and public health research. SACIDS-ACE provides research equipment to some institutions e.g. universities. AISA runs community outreach programmes to provide research resources to underprivileged schools. ITOCA assists institutions in developing ICT programmes and provides technical support to implement these programmes.

#### 3.2.5. Increasing research awareness

One organisation had as its objective to increase research awareness in Africa. AISA aims to promote knowledge creation as a fundamental aspect of development in Africa by encouraging research as a career choice for young people, as they leave school.

#### 3.2.6. Research collaboration

All RCS organisations in this study collaborated with other organisations – both in and out of Africa, for research. Some examples of intra-African collaborations included: AFIDEP, partnering with other training institutions and research organisations in Africa for research training workshops, and providing knowledge transfer platforms during these events. CRENC collaborates with other institutions such as universities, pharmaceutical, biotechnology and medical industries to provide technical assistance on clinical research projects. AISA collaborates with African multilateral organisations and provides research-based policy advice.

### 3.3. Analysis of RCS initiatives and activities using CORE framework

Table 3 presents the CORE Africa framework describing relevant factors that influence research capacity in Africa [18]. Figure 2 shows how many RCS organisations or initiatives identified in this study addressed each of the factors outlined in table 3. Priority levels were defined as “high priority” if a factor was being addressed by 65% or more of the identified RCS organisations; “medium priority” if addressed by 40-65% of the RCS organisations; “low priority” if addressed by 15-39% and “critical” if there were less than 15% of RCS organisations working on that factor.

**Figure 2:**
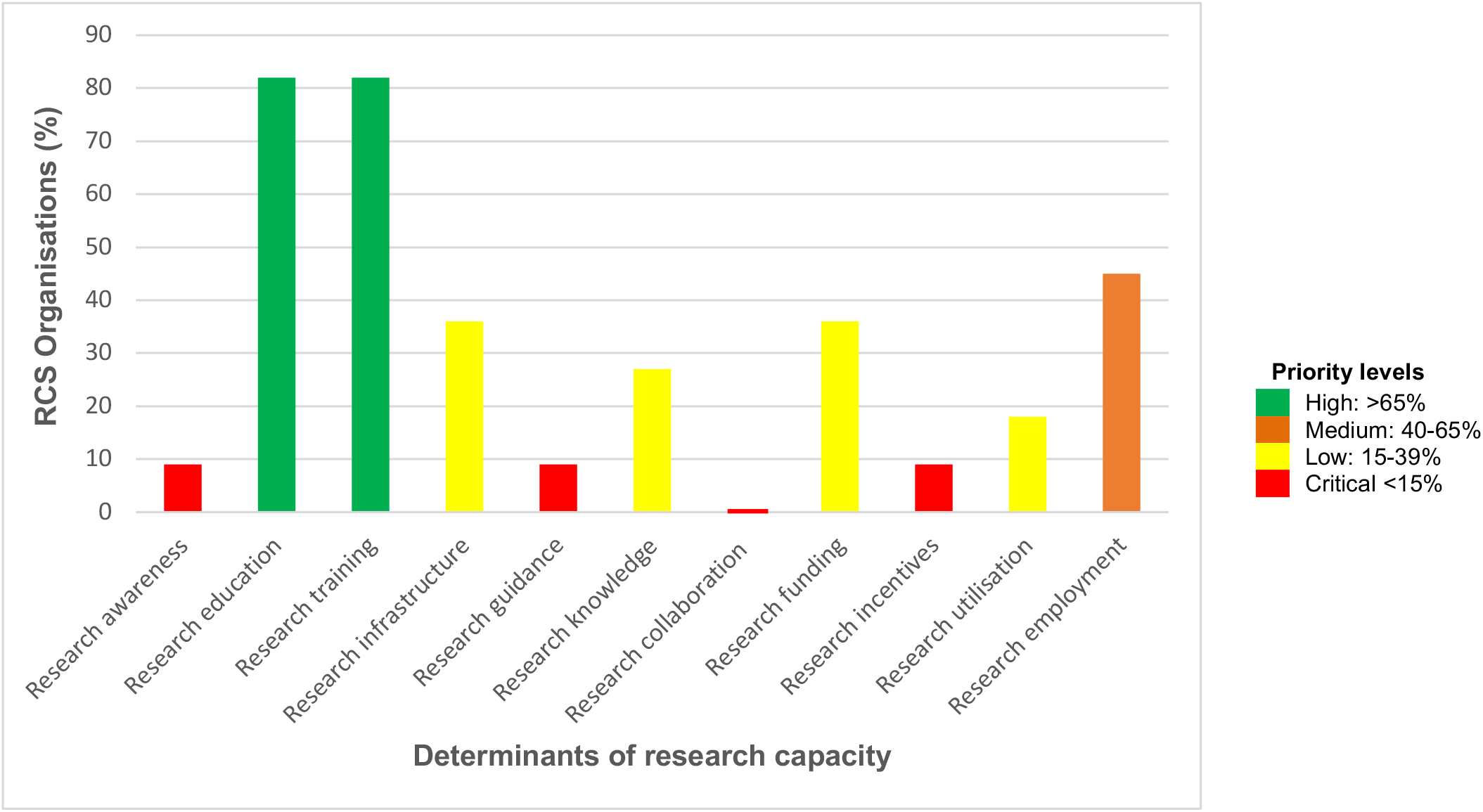
Factors addressed by RCS organisations in Africa.

**Table 3:** Relevant factors that influence research capacity in Africa

| Factor | Description |
| --- | --- |
| Research awareness | Increasing public understanding on the need for and importance of research, and encouraging engagement in research activity |
| Research education | Providing research knowledge through academic programs |
| Research training | Providing opportunities for research skill development |
| Research infrastructure | Providing resources, facilities and equipment required for producing and disseminating research e.g. computers, internet facilities and lab equipment |
| Research guidance | Providing leadership, counselling and supervision for research e.g. through mentorship |
| Research knowledge | Producing and publishing research findings |
| Research collaboration | Working with others to produce research and creating initiatives to help promote collaboration among African researchers |
| Research funding | Providing funds for research |
| Research incentives | Activities that encourage or motivate researchers and acknowledge research activity/output e.g. rewarding innovative projects, giving prizes, fee waivers, bonuses etc |
| Research utilisation | Using research findings for program development and policy-making |
| Research employment | Providing job opportunities and career-routes for researchers |

Two factors emerged as high priorities. These were providing research education and providing research training opportunities. 82% of the identified RCS organisations had programs or initiatives addressing these two areas.

One medium priority factor was found, which was providing job opportunities and career routes for African researchers. 45% of the RCS organisations had job opportunities for researchers.

Providing research resources and equipment, producing research knowledge, providing funding and facilitating the utilisation of research findings were seen to be low priorities; while increasing research awareness, providing research guidance, providing incentives for research and promoting research collaboration were critical.

Table 4 shows the different target groups identified to be important populations that influence research capacity in Africa. Figure 2 shows which population groups were targeted by the RCS organisations identified in this study. Priority levels for population groups were defined using the same criteria described above – high (>65%); medium (40-65%); low (15-30%) and critical (<15%). Researchers were found to be a high-priority group for RCS in this study, targeted by a majority (73%) of the identified RCS organisations. Academics, research institutions and policy makers were all medium priorities, targeted by 66%, 46% and 46% of the RCS organisations, respectively. No organisations were found targeting research funders or the general public.

**Table 4:** Key target groups for RCS in Africa

| Population group | Individuals |
| --- | --- |
| Academics | Students, graduates |
| Researchers | Independent researchers, research groups, scientists |
| Research institutions | Universities, organisations, institutions, collaborators |
| Policy makers | Governments, decision-makers at different levels |
| Funders | Any funding bodies |
| General public | General public |

**Figure 3:**
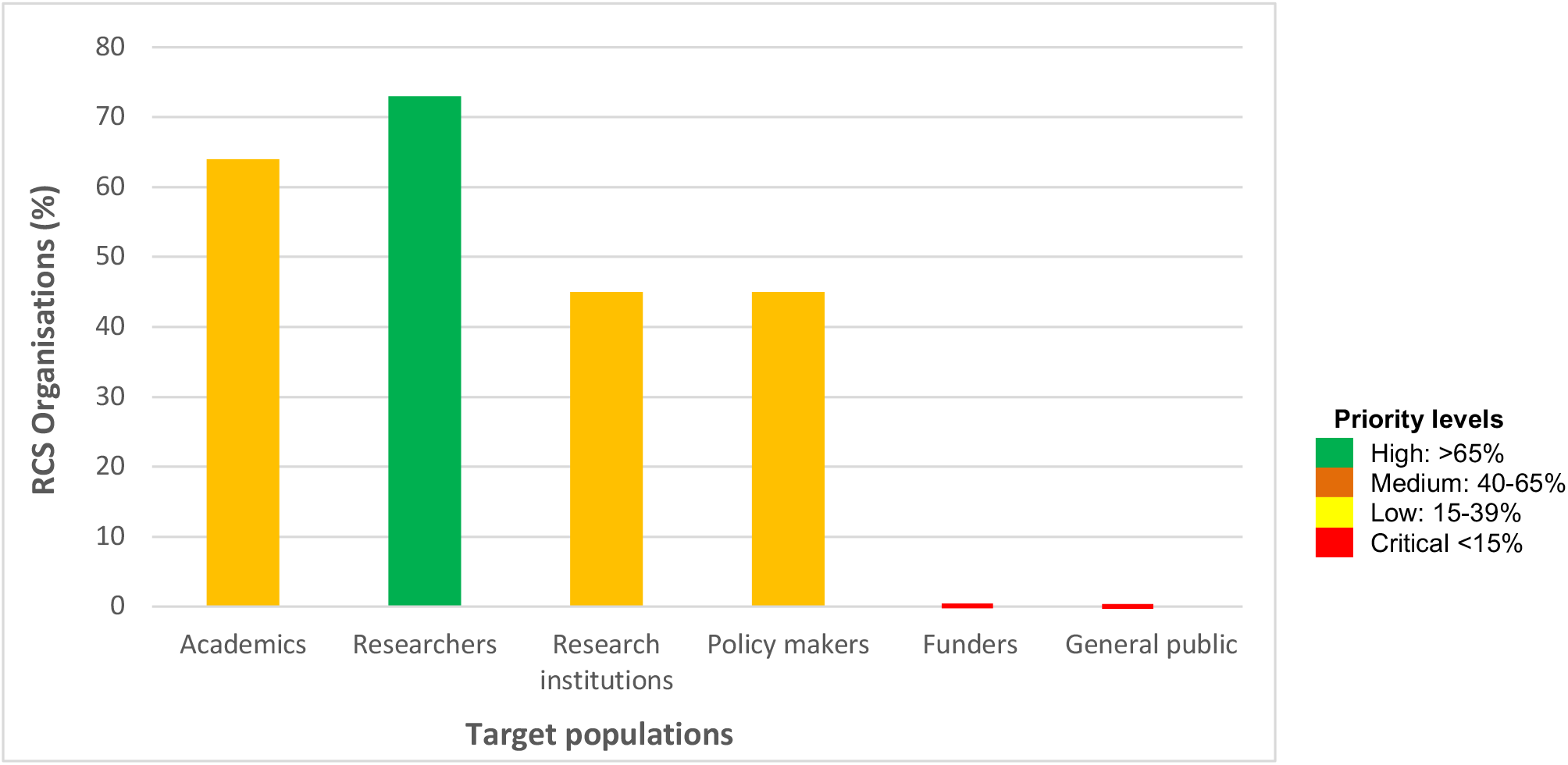
Population groups targeted by RCS organisations in Africa.

## Discussion

This study identified African-led organisations and initiatives that aim to strengthen research capacity in Africa. The study further explored what aspects of research capacity are considered priorities by RCS organisations in Africa, as well as those with little involvement. A useful feature of the framework used to assess the RCS organisations in this study is the fact that it breaks down the determinants of research capacity into relatable and easy-to-understand fragments, which together play a big role on research capacity at all levels (individual, organisational and national).

Most RCS organisations focused on providing opportunities for academic research and research training. Although research education was seen to be a priority in RCS, education systems in most African countries are not research-oriented [20]. Research is often introduced very late in the academic curriculum, mostly after secondary education. As such, academics at secondary school level, including teachers, school administrators and teacher trainers, have little or no involvement in research. At university level where research is introduced, academic curricula often have a poor research focus and fail to emphasise the need for research [21]. Most African universities aim to teach and confer degrees, unlike universities in other parts of the world which use new teaching models that focus on innovation [22]. Many African universities also do not incorporate new research modalities into established academic curricula, hence, producing graduates with worldviews and skill sets not suited to current world needs [23]. The education system not being research-oriented inevitably leads to poor quality research and a low research output.

The second priority area for RCS identified was the provision of research training opportunities. However, this finding stands in contrast to evidence from other studies reporting a lack of research training facilities as a key barrier to doing research in Africa [16, 26]. Academic institutions such as universities are usually the primary providers of research training opportunities in Africa. Meanwhile, most of these institutions lack facilities that enable the development of practical research skills and experience, such as well-equipped laboratories, computers, libraries and trained personnel [20, 24, 25].

The low priority factors identified in this study (providing research resources and equipment, producing research knowledge and providing funding for research), as well as the critical factors (increasing research awareness, providing research guidance, providing incentives for research and promoting research collaboration) support findings from existing literature, showing that these are some of the key challenges of doing research in Africa [16, 26, 27]. Research collaboration in Africa is mostly with researchers from developed countries such as the United States, the United Kingdom and France, with very little collaboration between African researchers [28]. Although the RCS organisations in this study were collaborating with other African organisations, there were no objectives or strategies aimed at encouraging or facilitating such collaboration. Intra-African research collaboration is key to pooling efforts towards a common goal and decreasing competition for funding, which is already limited.

The two population groups not targeted by RCS organisations in this study were funders and the general public. With the complexities of measuring the impact of capacity building programs in Africa, it is difficult for funders to measure, assess and account for investments in RCS initiatives [29]. As funders usually do not have direct access to local communities, they mostly rely on available evidence on program outcomes, effectiveness and gaps, which are then used to determine the need for funding and further investments [30]. Targeting funders in the RCS process and bringing this information to them is therefore key to establishing a negotiating position, vis-a-vis funding for RCS in Africa.

The overall aim of RCS is to be able to address challenges that affect the general population. It is therefore important for these individuals to understand how and why this is necessary. Public understanding of the need for research can help generate interest in research, improve research literacy and influence the use of research knowledge. Such understanding could also help create new funding streams to support research in Africa. For example, families with members suffering from chronic illnesses could be influenced through their understanding about research, to pool funds for research towards a particular disease. This can also be possible with philanthropists or other groups of people who are made to understand that research can help identify treatment approaches or improve health in local communities.

### Study Limitations

While multiple sources were searched in this study, we acknowledge that there are possibly several other RCS organisations and initiatives without an online presence, which could not be identified in this study. Cost barriers also did not allow for country visits and more detailed studies. The list of RCS organisations presented in this study are therefore non-exhaustive, and only intended to give a broad representation of RCS activities in Africa.

## Conclusion

This study highlights the importance of local leadership in the RCS process, and outlines the different approaches used by RCS organisations to improve research capacity in Africa. Our findings reveal gaps that could potentially influence research capacity in Africa, if addressed by future RCS programs or initiatives. Our concluding remarks and recommendations focus on five key areas:

- African researchers need to take ownership of the RCS process, in order to develop programs that are relevant to the African context and address priority needs in African countries
- Future RCS initiatives should address the factors that emerged as low priorities or critical factors in this study
- Education systems in Africa need to be more research-focused. Research needs to be introduced at an earlier stage of academic curricula, to help develop a research-oriented learning culture and promote interest in research
- RCS organisations in Africa need to develop initiatives that help funders understand what works to improve research capacity in Africa, to inform funding decisions for effective and sustainable programs
- African RCS organisations need to raise societal awareness on the importance of research in Africa, to enable appreciation for its need and create opportunities for engagement in research by the general public

## Acknowledgement

We would like to acknowledge the all researchers and panel members for their contribution in the findings of this report. We also acknowledge the CORE Africa research assistants and technical staff for their role in putting the data together.

## Appendix A Website links for RCS Organisations used in this study

APHRC

http://aphrc.org/our-work/research-strengthening

AFIDEP

https://www.afidep.org/about-us/how-we-work/capacity-strengthening/

REPOA

http://www.repoa.or.tz/about-us/

ITOCA

http://www.itoca.org

AISA

http://www.ai.org.za

AESA

https://aesa.ac.ke

SACORE

https://www.ncbi.nlm.nih.gov/pmc/articles/PMC4894455/#R5

ISHRECA

https://www.who.int/tdr/partnerships/initiatives/ishreca/en/

CARTA

http://cartafrica.org

SACIDS-ACE

http://www.sacids.org/sacids-ace-scholarship/

CRENC

https://crenc.org

